# Scaling patterns of body plans differ among squirrel ecotypes

**DOI:** 10.1101/2022.10.09.511490

**Authors:** Tate J. Linden, Abigail E. Burtner, Johannah Rickman, Annika McFeely, Sharlene E. Santana, Chris J. Law

## Abstract

Body size is often hypothesized to facilitate or constrain morphological diversity in the cranial, appendicular, and axial skeletons. However, how overall body shape scales with body size (i.e., body shape allometry) and whether these scaling patterns differ between ecological groups remains poorly investigated. Here, we test whether and how the relationships between body shape, body size, and limb lengths differ among species with different locomotory specializations, and describe the underlying morphological components that contribute to body shape variation among squirrel (Sciuridae) ecotypes. We quantified the body size and shape of 87 squirrel species from osteological specimens held at museum collections. Using phylogenetic comparative methods on these data, we found that 1) body shape and its underlying morphological components scale allometrically with body size, but these allometric patterns differ among squirrel ecotypes; 2) only ground squirrels exhibit a relationship between forelimb length and body shape, where more elongate species exhibit relatively shorter forelimbs; and 3) the relative length of the ribs and elongation or shortening of the thoracic region contributes the most to body shape variation across squirrels. Overall, our work contributes to the growing understanding of mammalian body shape evolution and how it is influenced by body size and locomotor ecology, in this case from robust subterranean to gracile gliding squirrels.

## Introduction

Body size is often hypothesized to be a line of least evolutionary resistance for morphological evolution (Marroig and Cheverud 2005), and evolutionary changes in body size have a strong influence on an organism’s ecological, physiological, morphological, and functional traits (Schmidt-Nielsen 1984; LaBarbera 1989; Calder 2001; Pryon and Burbink 2009). Because traits often scale with size, changes in body size facilitates the evolution of new traits that enable species to adapt to novel environments. However, extrinsic and intrinsic factors often constrain bodies towards certain sizes; therefore, in instances when evolutionary change in body size is limited, new adaptations can arise through evolutionary changes in the shape or proportions of traits (Zelditch et al. 2017). Unsurprisingly, a plethora of work has found that ecological factors affect the evolution of the shape and proportions of the skull (Janis 1990; Olsen 2017; Law et al. 2018; Arbour et al. 2019; Grossnickle 2020; Paluh et al. 2020), limbs (Van Valkenburgh 1985; Higham et al. 2015; Citadini et al. 2018; Baeckens et al. 2020) and vertebrae (Buchholtz 1998; Randau et al. 2016; Jones et al. 2018; Gillet et al. 2019; Luger et al. 2019; Adler et al. 2022). The evolution of diverse overall body shapes can also facilitate morphological, functional, and ecological innovations that can lead to increased diversification and niche specialization (Wiens et al. 2006; Collar et al. 2016; Law 2019; Friedman et al. 2020; Morinaga and Bergmann 2020).

Although the morphological patterns of body shape evolution are well-studied in vertebrates, including squamates (Wiens and Slingluff 2001; Grinham and Norman 2020; Bergman *et al.* 2020), fishes (Ward and Mehta 2010; Mehta *et al.* 2010; Friedman *et al.* 2019, Strauss 1985), and, more recently, carnivoran mammals (Law et al. 2019; Law 2021a), few have investigated evolutionary allometry between body shape and size. In fishes, body size explains 3–50% of body shape variation depending on taxonomic families, and larger fishes tend to exhibit more elongate bodies (Friedman *et al.* 2019). Similarly, in carnivoran mammals, allometric effects of skeletal size influence body shape variation (Law 2021a). However, the boundary between terrestrial and aquatic habitats affects these allometric patterns: terrestrial carnivorans tend to evolve more robust bodies with increasing size whereas aquatic carnivorans tend to evolve more elongate bodies with increasing size (Law 2021b), a congruent pattern with fishes (Friedman *et al.* 2019). This suggests that body shape allometries differ between locomotory ecologies. Associated with increasingly elongate body shapes is fin and limb size reduction (Gans 1975; Wake 1991; Ward and Mehta 2010; Wiens and Slingluff 2001; Skinner *et al.* 2008). In tetrapods, researchers have found that the forelimbs are usually reduced or lost prior to the hind limbs through evolutionary time (Wiens and Slingluff 2001; Gans 1975; Brandley et al. 2008; Morinaga and Bergmann 2017; Law et al. 2019). How locomotory ecologies affect relationships between body shape and limb lengths in mammals remains to be tested.

Despite observed convergence in body plans (e.g., Brandley et al. 2008; Friedman et al. 2016; Morinaga and Bergmann 2017; Bergmann and Morinaga 2019; Law 2022), similar body shapes can evolve through multiple pathways including elongation of the head, reduction of body depth, and elongation of the body axis via changes in total vertebral number and/or elongation of individual vertebrae (Parra-Olea and Wake 2001; Ward and Brainerd 2007; Ward and Mehta 2010; Collar et al. 2013; Law 2022). Because vertebral number is constrained in most mammals (Narita and Kuratani 2005), they evolve more elongate or robust bodies through changes in body depth and/or elongation or shortening of the skull and vertebrae rather than vertebral number. For example, the elongated neck found in giraffes is due to elongation of the cervical vertebrae rather than increase in the number of vertebrae (Badlangana 2009, Danowitz and Domalski 2015, Danowitz *et al.* 2015). In carnivorans, the elongation or shortening of the thoracolumbar regions and changes in rib lengths contribute most to variation in body shape, ranging from elongate weasels to robust bears (Law 2021a). Whether these patterns are similarly found in other mammalian clades is not known.

In this study, we used squirrels (family Sciuridae) as a model system to examine the effects of body size on body shape evolution. We also investigated the relationship between body shape and limb length as well as the underlying morphological components that contribute to body shape variation. Squirrels are qualitatively diverse, with body sizes ranging from 32 g least chipmunks to 8 kg Olympic marmots and body shapes ranging from the rotund bodies of marmots to the lithe bodies of gliding squirrels. In addition to their diverse body plans, squirrels exhibit varied locomotor ecologies and habitat use, including four ecotypes: ground squirrels that dig, tree squirrels that climb, gliding squirrels that glide between trees, and more versatile chipmunks that can dig and climb. Therefore, we also examined how morphological patterns differ between these four squirrel ecotypes.

Our objectives were three-fold. First, we examined the relationships between body size and body shape variation and between body shape variation and limb lengths. Second, we tested if these relationships differed between ecotypes. We predicted that ground squirrels would exhibit more robust bodies with increasing skeletal size and relatively shorter limbs. These phenotypes would provide more structural support and force production when digging large tunnel systems. We predicted that all other ecotypes (i.e., chipmunks, tree squirrels, and gliding squirrels) would evolve more elongate bodies with increasing skeletal size. Elongate bodies could facilitate heightened flexibility and maneuverability for large squirrels when navigating complex microhabitats such as tree branches. While it is known that the forelimbs of gliding squirrels are relatively longer than those of other ecotypes (Thorington and Heaney 1981; Peterka 1936; Bryant 1945; Grossnickle et al. 2020), the relationship between body shape and forelimb length remain unstudied. Accordingly, we predicted that gliding squirrels would exhibit relatively longer forelimbs with increasing body elongation to increase patagium surface area for gliding. In contrast, we predicted that chipmunks and tree squirrels would exhibit relatively shorter forelimbs with increasing body elongation following similar patterns found in terrestrial carnivoran mammals (Law et al. 2019). For our final objective, we examined which cranial and axial components contributed the most to overall body shape variation across squirrels and within each ecotype. We predicted that thoracolumbar elongation or shortening will be the biggest contributor to body shape variation, as this region provides the body’s primary structural support against gravity (Kardong 2014).

## Methods

### Quantifying squirrel body shape

We quantified squirrel body shape using 220 osteological specimens across 87 species. These specimens were sourced from the collections of 11 museums (Table S1). We used female, male, and sex-unknown individuals for our measurements in order to achieve the largest sample size possible per species and across species. Additionally, each specimen measured was fully mature, which we determined by verifying that the cranial exoccipital-basioccipital and basisphenoid-basioccipital sutures were fused and that all vertebrae and limb bones were ossified.

We used the head-body elongation ratio (hbER) to quantify body shapes, which was calculated as the sum of head length (L_H_) and body length (L_B_) divided by the body depth (L_R_): hbER = (L_H_ + L_B_)/L_R_ (Fig. 1). We measured head length as the condylobasal length of the cranium from the anteriormost point on the premallixa to the posteriormost point on the surfaces of the occipital condyles. We estimated body length by summing the centrum lengths (measured along the ventral surface of the vertebral centrum) of each cervical, thoracic, lumbar, and sacral vertebrae. All linear measurements were taken to the nearest 0.01 mm using digital calipers. We estimated body depth as the average length of the four longest ribs. Each rib was measured as a curve from the end of the capitulum to the point of articulation with the costal cartilage using a flexible measuring tape.

**Fig. 1.**
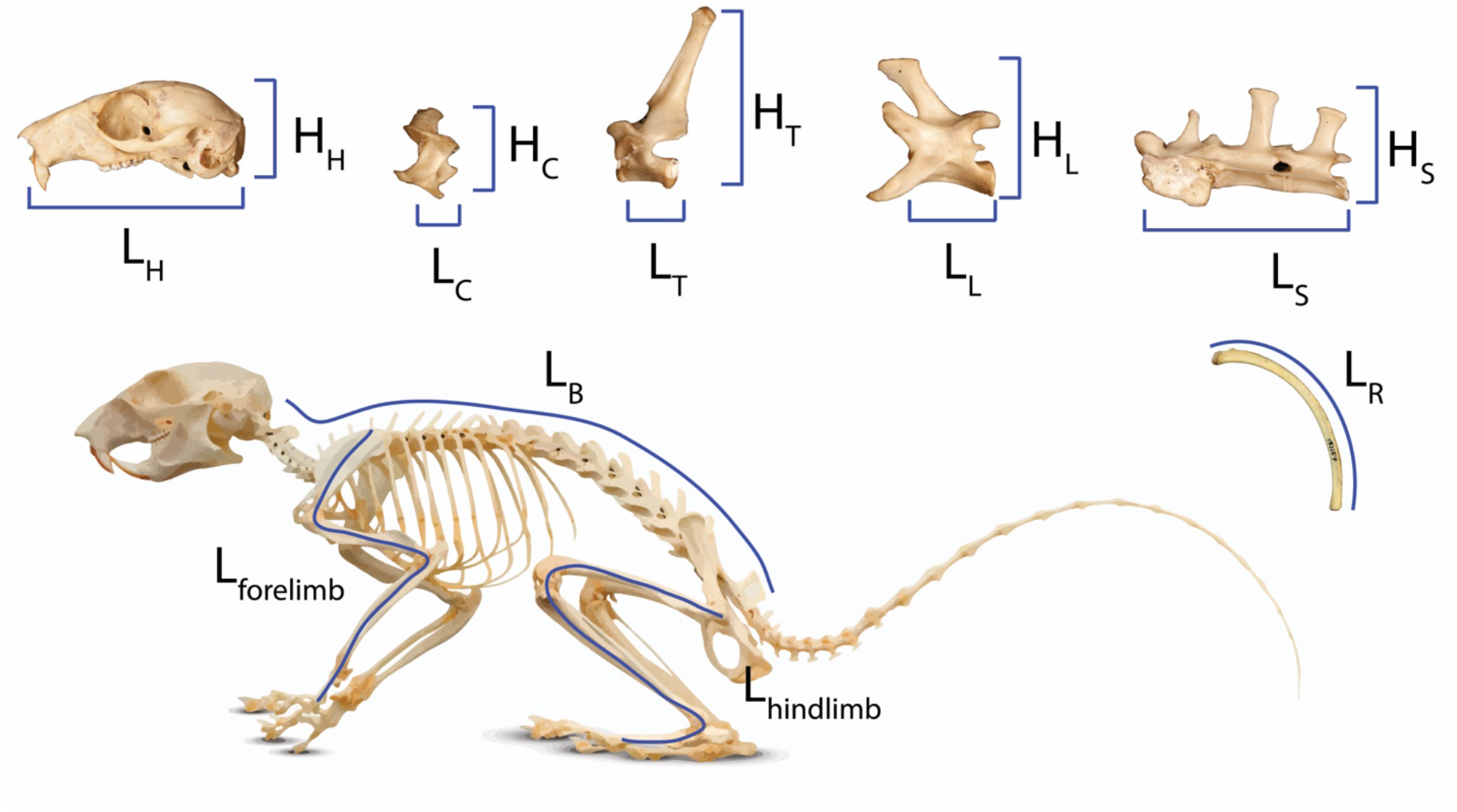
Measurements of body regions used to calculate head-body elongation ratio (hbER), head ER, and axial elongation index (AEI) of the cervical, thoracic, lumbar, and sacral regions. L_x_ = lengths and H_x_ = heights. hbER = (L_H_ + L_B_)/Lr AEI_v_ =∑L_v_/mean(H_v_). H = head; R = rib; C = cervical; T = thoracic; L =lumbar; S = sacral; B = body length.

We also quantified the underlying cranial and axial components that contribute to body shape variation. Head elongation ratio (head ER) was calculated by dividing cranial length (L_H_) by cranial height (H_H_). We used a modified version of the axial elongation index (AEI; Ward and Brainerd 2007; Law et al. 2019) to examine how each vertebral region (i.e., cervical, thoracic, lumbar, and sacral) contributes to body shape variation. For each vertebral region (V), we calculated AEI_V_ as the total sum of vertebral lengths (L_V_ measured along the ventral surface of the vertebral centrum) divided by the average vertebral height (H_V_; measured from the ventral surface of the centrum to the tip of the neural spine): AEI_V_ = ∑L_V_/mean(H_V_). We quantified skeletal size using the geometric mean of our linear measurements (Nth root of the product of our measurements for the cranium, vertebrae, and ribs; N = 11) (Mosimann 1970; Klingenberg 2016).

Lastly, we measured the lengths of the forelimb and hind limb. Forelimb length was estimated by summing the lengths of the scapula (from the dorsalmost point on the glenoid fossa to the ventralmost point on the inferior angle), humerus (from the dorsalmost point on the humeral head to the ventralmost point on the capitulum), radius (from the dorsalmost point on the radial head to the ventralmost point on the styloid process), and the third metatarsals (from the distalmost point on the metatarsal head to the proximalmost point on the metatarsal base). Hind limb length was estimated by summing the lengths of the femur (from the dorsalmost point on the femoral neck to the ventralmost point on the patellar surface), tibia (from the dorsalmost point on the intercondylar eminence to the ventralmost point on the articular surface), and third metacarpal (from the distalmost point on the metacarpal head to the proximalmost point on the metacarpal base). To increase species sample sizes, we used reduced limb datasets in which total limb lengths consisted of just the long bones (i.e., humerus and radius for the forelimb and femur and tibia for the hind limb). Our findings with these datasets indicated that the major patterns remain the same (see Results).

### Ecotype data

We categorized species into four ecotypes: chipmunks (*n* = 15), gliding squirrels (*n* = 11), ground squirrels (*n* = 29), and tree squirrels (*n* = 32). We categorized squirrels based on locomotion and nest location using natural history information from the *Handbook of the Mammals of the World* (Wilson and Mittermeier 2009) and the Animal Diversity Web (https://animaldiversity.org/). We classified tree squirrels as species that nest in trees and display both arboreal and scansorial locomotion, gliding squirrels as species with derived morphologies (i.e., patagia) for gliding locomotion, and ground squirrels as species that nest in underground burrows and display fossorial locomotion (Hayssen 2008). Our fourth ecotype group was chipmunks (genus *Tamias*), which display the broadest range of locomotor and nesting behaviors; species are considered terrestrial, semi-fossorial, or semi-arboreal depending on the source consulted, but none are considered fully fossorial or arboreal.

### Statistical Analyses

All analyses were performed under a phylogenetic framework using Upham *et al*’s (2019) recent phylogeny of mammals pruned to include just the 87 studied squirrels. We took the natural logarithm of all traits prior to statistical analyses and performed all analyses in R (2022).

We tested for allometric relationships between body shape and body size using a phylogenetic generalized least squares (PGLS) regression. We then tested if body shape allometry differed among ecotypes using PGLS regressions with an ANCOVA design. To determine if there was significant body shape allometry, we generated 95% confidence intervals using bootstrapping (1000 replications). Confidence intervals that deviated from an isometric slope of 0 were interpreted as exhibiting significant positive allometry (slope > 0) or negative allometry (slope < 0). The isometric slope was set as 0 because hbER is a dimensionless ratio. Additionally, we determined whether the allometric slopes differed between ecotypes by comparing the mean slope of each ecotype with the 95% confidence intervals of the mean slope of the other ecotypes. All regression coefficients were simultaneously estimated with phylogenetic signal in the residual error as Pagel’s lambda using the R package phylolm v2.6.2 (Tung Ho and Ané 2014).

We examined if the relationships between limb length and body shape using PGLS and tested if these relationships differed among ecotypes using PGLS regressions with an ANCOVA design in phylolm. We excluded 11 species from which we were unable to collect limb data. We determined if each ecotype exhibited a significant limb~body shape relationship based on whether the 95% confidence intervals from 1000 bootstrap replications deviated from an isometric slope of 0. We size-corrected limb lengths prior to analyses by extracting residuals for each trait against the geometric mean using a PGLS. We also tested if size-corrected limb lengths differed among ecotypes. Differences among ecotypes were considered significant in each measure of limb length if an ecotype’s value for mean limb length was outside of other ecotypes’ 95% bootstrap confidence intervals for that measure of limb length. We also ran PGLS regressions on the full limb datasets that included the metacarpals and metatarsals.

Lastly, to determine which morphological components (i.e., cranium, ribs, cervical vertebrae, thoracic vertebrae, lumbar vertebrae, and sacrum) contributed the most to variation in body shape, we performed phylogenetic multiple regressions using the R package RRPP v1.0.0 (Collyer and Adams 2018). Statistical significance was determined using the random residual permutation procedure (RRPP) with 1000 iterations (Adams and Collyer 2018). We performed phylogenetic multiple regressions for the whole clade as well as each of the four ecotypes.

## Results

### Body shape allometry

Across all squirrels, we found that there was no significant relationship between body size and body shape (PGLS: R^2^ = 0.01, λ = 0.80, slope [95% CI] = −0.02 [−0.07:0.02]; Table S2). However, including ecotype*size as an interaction term indicated that allometric trends in body shape differed between ecotypes (PGLS: R^2^ = 0.52, λ = 0.00; Fig. 2). Gliding squirrels (0.12 [0.06:0.19]) and chipmunks (0.24 [0.03:0.46]) exhibited positive allometry, indicating that gliding and chipmunk species evolved more elongate bodies with increasing skeletal size. In contrast, ground squirrels (−0.11 [−0.15:−0.06]) exhibited negative allometry, indicating that ground squirrels evolved more robust bodies with increasing skeletal size. Tree squirrels did not exhibit a significant relationship between body shape and skeletal size (−0.02 [−0.08:0.03]).

**Fig. 2.**
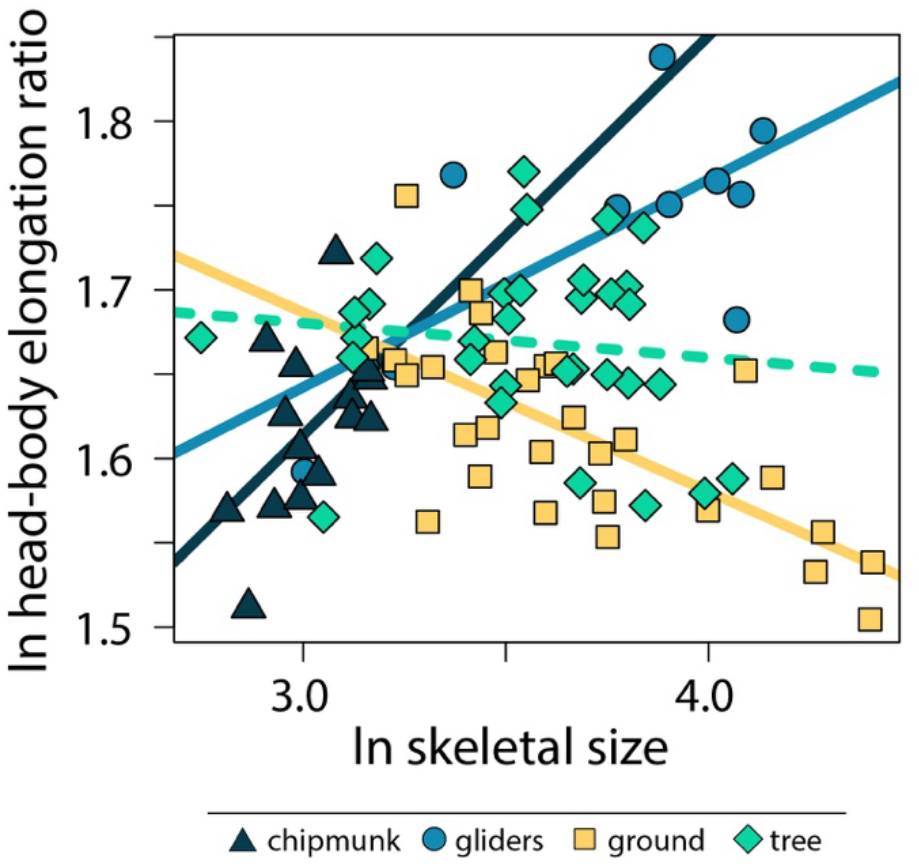
Scatter plot of ln skeletal size and ln head-body elongation ratio (hbER). Relationships between skeletal size and hbER were tested using PGLS with an ANCOVA design. Solid lines indicate significant relationships, and dashed lines indicate non-significant relationships. Confidence intervals that deviated from an isometric slope of 0 were interpreted as exhibiting significant positive allometry (slope > 0) or negative allometry (slope < 0).

Within the body shape components, head ER (0.18 [0.13:0.23] and cervical AEI (0.22 [0.18:0.26]) scaled with positive allometry with body size whereas lumbar ER (−0.09 [−0.16:0.02]) scaled with negative allometry across all squirrels. These results indicate that squirrels exhibit more elongate heads and cervical regions but more robust lumbar regions with increasing skeletal size. In contrast, thoracic and sacral regions and size-corrected rib length did not scale with skeltal size (Table S2).

Allometric trends in body shape components also differed among ecotypes. For gliding squirrels, we found positive allometry for ln head ER (0.21 [0.12 : 0.30]), ln cervical AEI (0.29 [0.20 : 0.37]), and ln lumbar AEI (0.15 [0.01 : 0.29]; Fig. 3; Table S2). Tree squirrels exhibited a positive slope for only ln cervical AEI (0.27 [0.20 : 0.35]). Ground squirrels showed negative allometry for ln thoracic AEI (−0.17 [−0.27 : −0.09]) and ln lumbar AEI (−0.23 [−0.32 : −0.14]) but positive allometry for ln cervical AEI (0.13 [0.07 : 0.18]), ln head ER (0.22 [0.15 : 0.29]), and ln size-corrected rib length (0.05 [0.00 : 0.08]). Chipmunks showed a positive trend for ln cervical AEI (0.50 [0.31 : 0.67]). We found no significant allometry for ln sacral AEI in any ecotype (Fig. 3; Table S2).

**Fig. 3.**
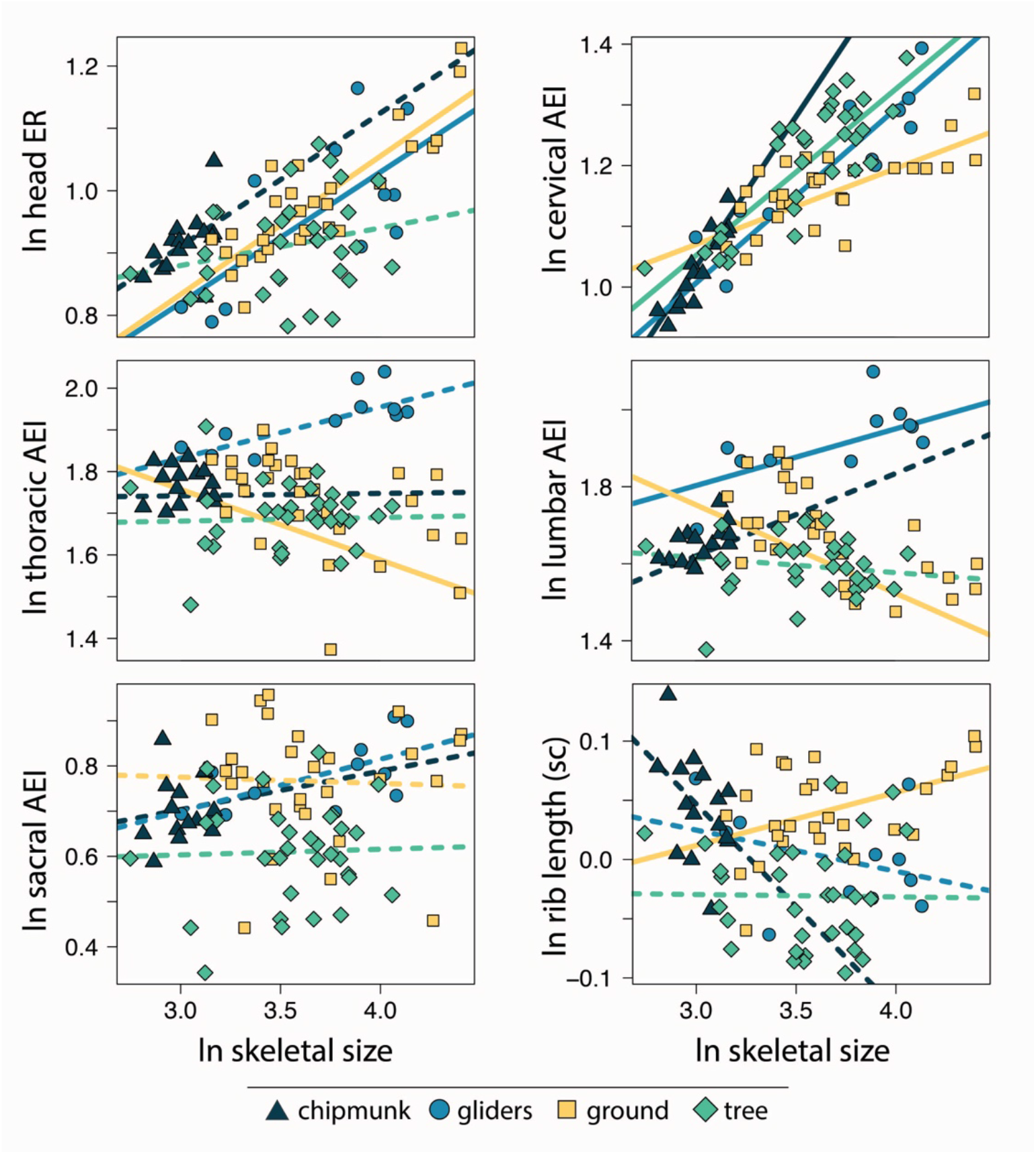
Scatter plot of ln skeletal size and ln each morphological component underlying body shape variation. Relationships between skeletal size and each morphological component were tested using PGLS with an ANCOVA design. Confidence intervals that deviated from an isometric slope of 0 were interpreted as exhibiting significant positive allometry (slope > 0) or negative allometry (slope < 0). Solid lines indicate significant relationships, and dashed lines indicate non-significant relationships.

### Relationships between body shape and limb length

Across all squirrels, we found that body shape did not scale with either size-corrected forelimb length (0.08 [−0.17:0.34]) or size-corrected hind limb length (0.08 [−0.17:0.29]; Table S3). Findings remained largely the same when using the full forelimb (scapula + humerus + radius + metacarpal) and hind limb (femur + tibia + metatarsal) datasets (Table S3).

Relationships between body shape and size-corrected forelimb lengths differed between ecotypes. Ground squirrels exhibited a negative relationship between size-corrected forelimb length and body shape (slope [95% CI] = −0.391 [−0.672 : −0.098]), whereas the remaining ecotypes did not exhibit significant relationships between relative forelimb length and body shape (chipmunk slope = −0.01 [−0.56:0.48]; gliding squirrel slope = 0.37 [−0.04:0.86]; tree squirrel slope = 0.01 [−0.32:0.32]). In contrast, none of the ecotypes exhibited significant relationships between relative hind limb length and body shape (chipmunk slope = −0.05 [−0.45:0.32]; gliding squirrel slope = 0.33 [−0.35:09]; ground squirrel slope = 0.02 [−0.31:0.34]; tree squirrel slope = −0.14 [−0.55:0.25]; Fig. 4).

**Fig. 4.**
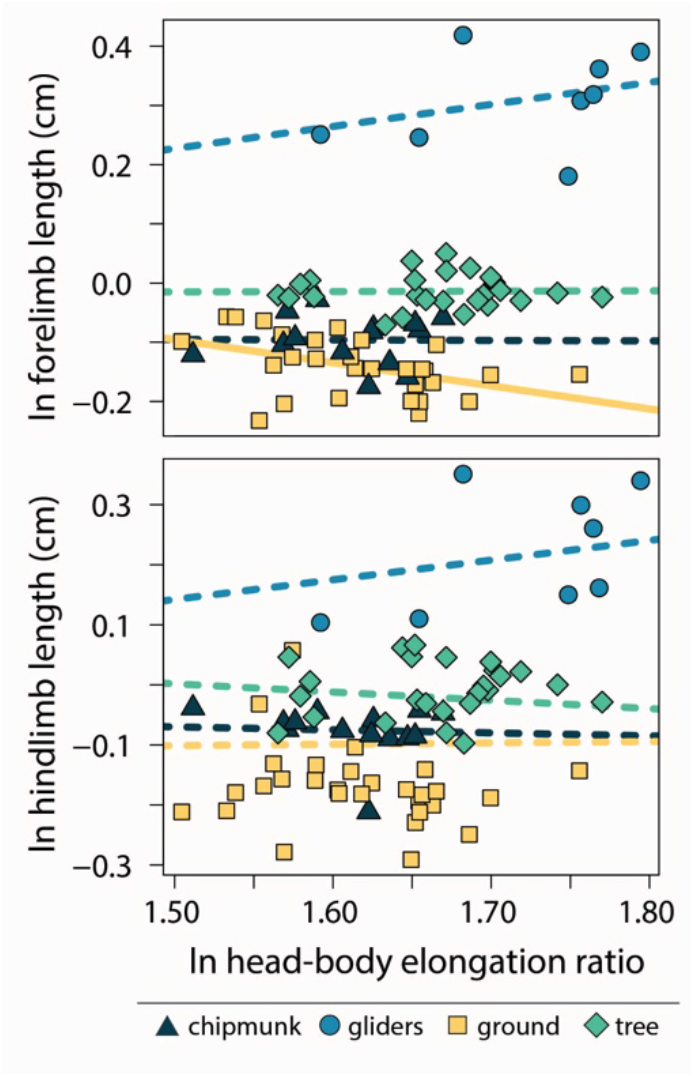
Scatter plot of ln head-body elongation ratio (hbER) and ln size-corrected forelimb and hind limb lengths. Relationships between hbER and limb lengths were tested using PGLS with an ANCOVA design.Confidence intervals that deviated from an isometric slope of 0 were interpreted as exhibiting significant positive relationship (slope > 0) or negative relationship (slope < 0). Solid lines indicate significant relationships, and dashed lines indicate non-significant relationships.

Gliding squirrels exhibited relatively longer forelimbs (residuals [95%CI] = 0.49 cm [0.17:0.80 cm]) than all other ecotypes (chipmunks = 0.07 cm [−0.23:0.36 cm]; ground squirrels = 0.03 cm [−0.27:0.32 cm]; tree squirrels = 0.16 cm [−1.45:0.46 cm]) (Fig. 5). Gliding squirrels also exhibited relatively longer hind limbs (0.23 cm [−0.17:0.58 cm]) than ground squirrels (−0.09 cm [−0.43:0.22 cm]) but not chipmunks (−0.07 cm [−0.42:0.26 cm]) or tree squirrels (−0.01 cm [−0.37:0.32 cm]; Fig. 5).

**Fig. 5.**
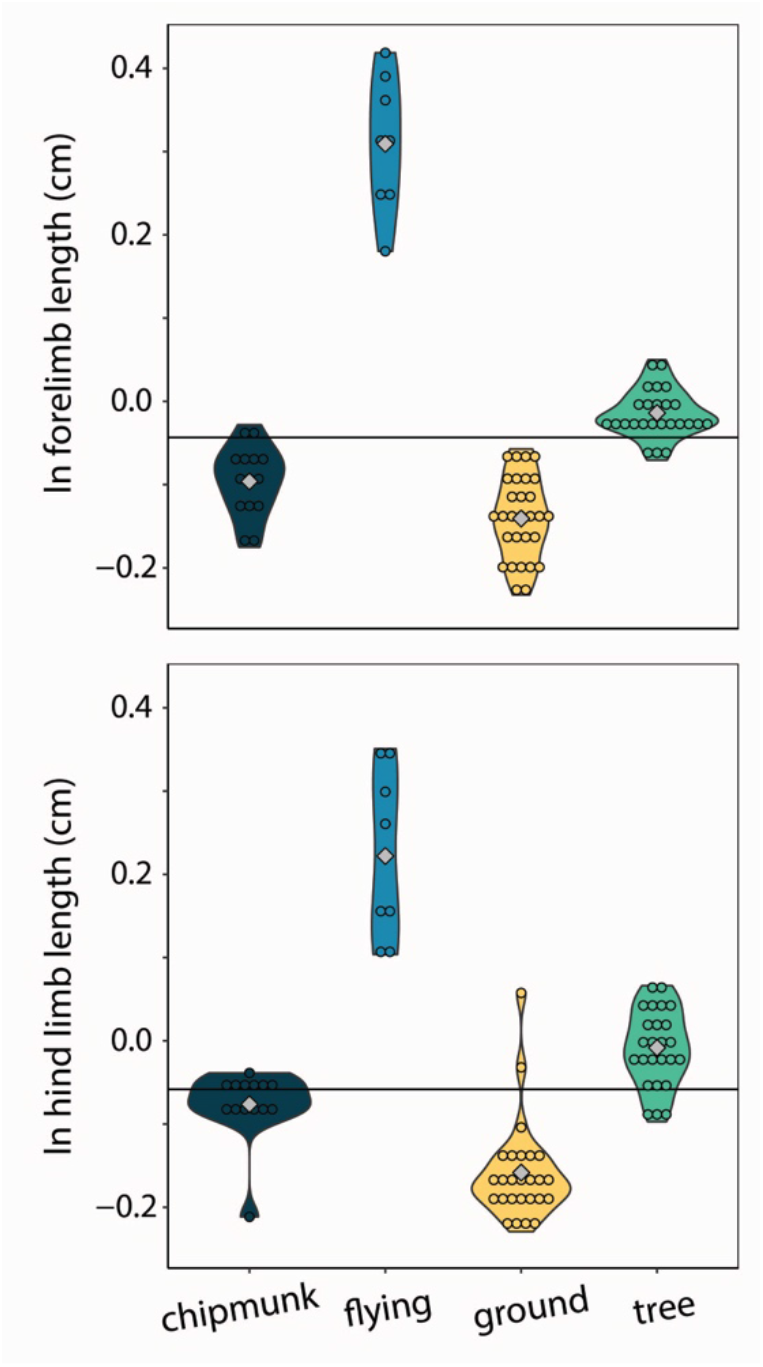
Violin plots of size-corrected forelimb length and size-corrected hind limb length. Gliding squirrels exhibited relatively longer forelimbs than all other ecotypes and relatively longer hind limbs than ground squirrels. The horizontal black line indicates the mean size-corrected limb length for all ecotypes.

### Predictors of body shape variation

We found that the relative length of the ribs (R^2^ = 0.64; P = 0.001) and elongation or shortening of the thoracic region (R^2^ = 0.19; P = 0.001) and sacral region (R^2^ = 0.11; P = 0.001) contributed the most to overall body shape variation across squirrels (Table 1). Elongation or shortening of the head, cervical, and lumbar regions contributed less than 2%. When examining each ecotype separately, we found that the relative length of the ribs was also the best predictor of body shape variation in chipmunks (R^2^ = 0.48; P = 0.001), ground squirrels (R^2^, = 0.34; P = 0.001), and tree squirrels (R^2^ = 0.54; P = 0.001). In contrast, elongation or shortening of the head (R^2^ = 0.55; P = 0.003) was the best predictor of body shape variation in gliding squirrels. Contributions to body shape variation from the remaining skeletal components ranged from 0 to 25% (Table 1).

**Table 1.** Results of the phylogenetic multiple regression with the random residual permutation procedure (RRPP) to determine which skeletal components contributed most to body shape variation across all squirrels and within each ecotype. Bold p-values indicate significance (α = 0.05).

| Skeletal Component | Df | SS | MS | R2 | F | Z | P |
| --- | --- | --- | --- | --- | --- | --- | --- |
| A. All ecotypes combined |  |  |  |  |  |  |  |
| ln head elongation ratio | 1 | 0.000 | 0.000 | 0.00 | 4.28 | 1.65 | **0.044** |
| ln cervical AEI | 1 | 0.000 | 0.000 | 0.00 | 0.15 | -0.60 | 0.721 |
| ln thoracic AEI | 1 | 0.011 | 0.011 | 0.19 | 312.24 | 7.98 | **0.001** |
| ln lumbar AEI | 1 | 0.001 | 0.001 | 0.02 | 30.24 | 3.81 | **0.001** |
| ln sacral AEI | 1 | 0.006 | 0.006 | 0.11 | 175.30 | 6.16 | **0.001** |
| ln size-corrected rib length | 1 | 0.038 | 0.038 | 0.64 | 1066.64 | 10.30 | **0.001** |
| Residuals | 80 | 0.003 | 0.000 | 0.05 |  |  |  |
| Total | 86 | 0.059 |  |  |  |  |  |
| B. Chipmunks |  |  |  |  |  |  |  |
| ln head elongation ratio | 1 | 0.002 | 0.002 | 0.08 | 167.98 | 5.21 | **0.001** |
| ln cervical AEI | 1 | 0.004 | 0.004 | 0.16 | 345.02 | 6.34 | **0.001** |
| ln thoracic AEI | 1 | 0.002 | 0.002 | 0.08 | 175.56 | 4.98 | **0.001** |
| ln lumbar AEI | 1 | 0.004 | 0.004 | 0.15 | 320.14 | 5.79 | **0.001** |
| ln sacral AEI | 1 | 0.001 | 0.001 | 0.04 | 86.53 | 4.32 | **0.001** |
| ln size-corrected rib length | 1 | 0.013 | 0.013 | 0.48 | 1031.09 | 8.31 | **0.001** |
| Residuals | 8 | 0.000 | 0.000 | 0.00 |  |  |  |
| Total | 14 | 0.027 |  |  |  |  |  |
| C. Gliding squirrels |  |  |  |  |  |  |  |
| ln head elongation ratio | 1 | 0.002 | 0.002 | 0.55 | 52.22 | 3.24 | **0.003** |
| ln cervical AEI | 1 | 0.000 | 0.000 | 0.00 | 0.30 | -0.24 | 0.604 |
| ln thoracic AEI | 1 | 0.000 | 0.000 | 0.00 | 0.35 | -0.15 | 0.551 |
| ln lumbar AEI | 1 | 0.001 | 0.001 | 0.25 | 23.71 | 2.50 | **0.004** |
| ln sacral AEI | 1 | 0.000 | 0.000 | 0.05 | 4.60 | 1.32 | 0.084 |
| ln size-corrected rib length | 1 | 0.000 | 0.000 | 0.11 | 10.57 | 1.77 | **0.031** |
| Residuals | 4 | 0.000 | 0.000 | 0.04 |  |  |  |
| Total | 10 | 0.004 |  |  |  |  |  |
| D. Ground squirrels |  |  |  |  |  |  |  |
| ln head elongation ratio | 1 | 0.004 | 0.004 | 0.24 | 60.07 | 4.59 | **0.001** |
| ln cervical AEI | 1 | 0.002 | 0.002 | 0.13 | 33.52 | 3.61 | **0.001** |
| ln thoracic AEI | 1 | 0.001 | 0.001 | 0.04 | 10.85 | 2.43 | **0.006** |
| ln lumbar AEI | 1 | 0.000 | 0.000 | 0.00 | 0.68 | 0.21 | 0.428 |
| ln sacral AEI | 1 | 0.003 | 0.003 | 0.16 | 40.08 | 3.77 | **0.001** |
| ln size-corrected rib length | 1 | 0.006 | 0.006 | 0.34 | 86.92 | 5.06 | **0.001** |
| Residuals | 22 | 0.001 | 0.000 | 0.09 |  |  |  |
| Total | 28 | 0.017 |  |  |  |  |  |
| E. Tree squirrels |  |  |  |  |  |  |  |
| ln head elongation ratio | 1 | 0.000 | 0.000 | 0.00 | 1.44 | 0.78 | 0.23 |
| ln cervical AEI | 1 | 0.002 | 0.002 | 0.16 | 91.96 | 4.78 | **0.001** |
| ln thoracic AEI | 1 | 0.002 | 0.002 | 0.23 | 130.62 | 5.30 | **0.001** |
| ln lumbar AEI | 1 | 0.000 | 0.000 | 0.01 | 7.48 | 2.06 | **0.013** |
| ln sacral AEI | 1 | 0.000 | 0.000 | 0.01 | 3.31 | 1.38 | 0.082 |
| ln size-corrected rib length | 1 | 0.006 | 0.006 | 0.54 | 305.28 | 7.86 | **0.001** |
| Residuals | 25 | 0.000 | 0.000 | 0.04 |  |  |  |
| Total | 31 | 0.011 |  |  |  |  |  |

## Discussion

### Body shape allometry and limb length evolution

Although size is known to influence variation in cranial, vertebral, and appendicular shape within and across species (e.g., Jones et al 2018; Baliga and Mehta 2016; Zelditch et al 2017; Stepanova and Womack 2020; Law et al 2022), few studies have tested allometric patterns in overall body shapes (but see Friedman *et al.* 2019; Law 2021a). Here, our results indicate that the relationship between body size and body shape is nuanced by ecological specialization; both body shape allometry and relationships between body shape and limb lengths differed between ecotypes (Fig. 2). Specifically, we found positive allometry in chipmunk and gliding squirrel body shapes, negative allometry in ground squirrel body shapes, and no significant effect of body size on the evolution of tree squirrel body shapes.

As predicted, ground squirrels exhibited more robust bodies and relatively shorter limbs with increasing skeletal size. These large, robust bodies along with relatively short forelimbs provide the necessary support, increased mechanical advantage, and force production (Lagaria and Youlatos 2006, Samuels and Van Valkenburgh 2008) needed to dig large burrow systems (Goldstein 1972, Casinos *et al.* 1993). This negative body shape allometry (i.e., evolution towards more robust bodies with increasing size) is consistent with what is found in terrestrial carnivorans (Law et al. 2019; Law 2021b). In large terrestrial mammals, robust body shapes (Law 2021a) and low spinal flexibility (i.e., dorsostability) of the vertebral column (Jones 2015a, Halpert *et al.* 1987) provide increased support against gravity for their heavier bodies (Kardong 2014). For example, the lumbar vertebrae of large bovids and felids have more robust centra than found in smaller species, providing more stability against sagittal bending during running (Jones 2015a). However, in Sciuridae, negative body shape allometry was only observed in ground squirrels, suggesting that fossoriality facilitated selection towards more robust bodies with increasing size.

Ground squirrels evolved relatively shorter forelimbs with increasing body elongation, reflecting the only significant trend observed between limb lengths and body shape in squirrels. The reduction or loss of limbs tend to evolve with body elongation in ectotherms (Gans 1975; Wake 1991; Wiens and Slingluff 2001; Skinner *et al.* 2008) and musteloid mammals (Law *et al.* 2019). With the notable exception of snakes (Gans 1975), this trend tends to be found in species that dig (Wake 1991; Lee 1998; Gans 1975 Rieppel 1988), spend significant time hunting in burrows (Law *et al.* 2019), or shelter in leaf litter and other surface debris (Gans 1975).

Additionally, elongate, limb-reduced bodies tend to be associated with a small body size (Lee 1998; Rieppel 1988). These trends are only corroborated by our results for ground squirrels, the most fossorial of the four ecotypes. This allometric trend may suggest that small, elongate ground squirrels rely more on their relatively shorter forelimbs to dig burrows compared to larger, more robust ground squirrels. Additionally, we found that the reduction of forelimbs in elongate species tended to evolve before the reduction of hind limbs, as seen in both mustelids (Law *et al.* 2019) and ectotherms (Wiens and Slingluff 2001; Gans 1975; Brandley et al. 2008; Morinaga and Bergmann 2017). Our research marked the first evidence of this trend in rodents.

In contrast to ground squirrels and terrestrial carnivorans, gliding squirrels exhibited positive body shape allometry, in which larger species exhibit more elongate bodies. Additionally, we confirm Thorington and Heaney’s (1981) hypothesis that small tree and gliding squirrels exhibit similarly robust bodies, while large gliding squirrels are more elongate than large tree squirrels. An elongate body could enable more aerodynamic and maneuverable gliding in large gliding squirrels, which face increased effects of gravity and drag due to their larger sizes. Because body mass is expected to increase proportionally to the cube of linear measurements, whereas wing area increases proportionally to the square, wing loading (body mass / patagium area) would naturally be higher in large gliding squirrels (Thorington and Heaney 1981). There are multiple pathways through which large gliding squirrels can compensate for this, including body elongation allometry (i.e., decreasing relative body weight with size) and limb length allometry (i.e., disproportionately increasing patagium area with size). The elongate forms of large gliding squirrels correspond to a 16% lower relative body weight than small gliding squirrels, decreasing the effects of wing loading on large squirrels (Thorington and Heaney 1981). Interestingly, relative limb length did not increase with body elongation (Fig. 4), possibly due to constraints from scansorial and gliding locomotion on forelimb length. Exceedingly long limbs could interfere with the squirrels’ ability to climb, reduce maneuverability when gliding, and risk more frequent breakage. Despite the lack of relationship between limb length and body elongation, gliding squirrels increase the area of their patagium through the styliform cartilage on their wrists (Johnson-Murray 1977; Thorington *et al.* 1998). The styliform extension, while isometric with respect to size (Thorington and Heaney 1981), could reduce selection towards positive allometry in limb length by both increasing maneuverability (acting similarly to a winglet on an airplane wing) and reducing wing loading (Thorington *et al.* 1998).

Despite the lack of relative limb length allometry, we confirm that gliding squirrels exhibited relatively longer forelimbs than all other ecotypes (Fig. 5; Thorington and Heaney 1981; Peterka 1936; Bryant 1945), a pattern that has been found across all gliding mammals (Grossnickle et al. 2020). Therefore, the elongate bodies and relatively longer forelimbs together in gliding squirrels could reduce the effects of wing loading on the gliding capabilities of larger species. Interestingly, the observation that gliders exhibited relatively longer hind limbs compared to the remaining ecotypes was not as well supported statistically (Fig. 5). A possible explanation for this pattern is that, because of the positioning of the forelimbs in the patagium, elongating the forelimbs may better support a wider plagiopatagium than elongating the hind limbs. Longer hind limbs would increase the uropatagium area and thus decrease the aspect ratio (wingspan^2^ / wing area) of gliding squirrels, which is already quite low compared to birds (Thorington and Heaney 1981). While a low aspect ratio could facilitate landing (Zimmerman 1932, Thorington and Heaney 1981) and increase maneuverability (Norberg 1995), Thorington and Heaney (1981) proposed that this low aspect ratio could decrease glide ratio (horizontal distance / altitude loss). A low glide ratio could negatively impact gliding ability, decreasing the selective pressure for long hind limb length in gliding squirrels. Furthermore, although longer hind limbs can increase patagium area, evolving relatively longer hind limbs risks increasing drag. Additionally, the relative hind limb lengths of gliding squirrels increased with body elongation when the metatarsals were included. The patagium connects at the ankle, so large gliders’ higher metatarsal length does not affect wing area despite uropatagium area being relatively larger in large gliding squirrels, possibly as an adaptation to compensate for their high wing loading (Thorington and Heaney 1981). This finding could instead be related to the start of gliding locomotion; larger, more elongate gliding squirrels may require more momentum from long metatarsals to propel themselves off of branches. Furthermore, aspects of the morphology other than forelimb and hind limb length, such as patagium musculature (Johnson-Murray 1977), could be altered to compensate for wing loading or increase maneuverability in large species.

Generalist chipmunks did not exhibit similar trends to fossorial ground squirrels; chipmunks showed neither a negative body shape allometric trend nor negative trend between relative limb length and body elongation. A possible explanation is that chipmunks exhibit a narrower range of body sizes, from the 32−50 g least chipmunk to the 66−150 g Eastern chipmunk (Roland 2009, Reid 2006, Kurta 1995). Chipmunks are also small compared to ground squirrels, which range from the 96–117 g white-tailed antelope squirrel to the 8 kg Olympic marmot. Chipmunks and small ground squirrels tend to burrow for shelter, whereas larger ground squirrels and marmots tend to build extensive burrow systems (Armstrong 2007). Therefore, these differences in size-related burrowing behavior and lack of body size diversity within chipmunks could explain their different trends compared to ground squirrels. Instead, chipmunks exhibit more elongate bodies with increasing skeletal size, an adaptation that may provide larger chipmunk species increased maneuverability when climbing trees and navigating tight tree hollows.

Lastly, there was no relationship between skeletal size and body shape in tree squirrels. An elongate body could be relatively unimportant compared to relative tail length for arboreal maneuvering. Tree squirrels exhibit relatively longer tails than ground squirrels (Hayssen 2008), which may facilitate balance during arboreal locomotion (Buck *et al.* 1925; Siegel 1970, Hayssen 2008) and help right falling squirrels (Fukushima et al. 2021). The relatively longer tails of larger squirrels could contribute to large tree squirrels’ ability to navigate arboreal terrain despite the lack of body shape allometry.

### Allometry of axial skeleton components

Locomotion can impact the evolution of the axial skeleton, especially the thoracolumbar region, which is the main axial region responsible for generating the propulsive forces necessary for locomotion (Kardong 2014; Boszczyk *et al.* 2001). Suspensory mammals with dorsostabile adaptations have higher variability in presacral vertebral number than running mammals with dorsomobile (high spinal flexibility) adaptations (Williams *et al.* 2019). As size increases, the energetic and biomechanical costs of dorsomobile running also increases. Therefore, a trade-off between stabilization and efficiency may have led to increased lumbar stability with increasing body size in large running mammals such as bovids (Jones 2015). Here, we found that allometry influenced the evolution of the thoracolumbar region in only the ground and gliding squirrels. Unsurprisingly, ground squirrels evolved a more robust thoracic region and relatively longer ribs with increasing skeletal size. A more robust thoracic region and rib cage contributes to dorsostability in large squirrels (Boszczyk *et al.* 2001; Jones 2015a, b), which could support their bodies as they dig large tunnel systems. In contrast, gliding squirrels evolved a more elongate lumbar region. The increased elongation in the lumbar region of larger gliding squirrels could be an adaptation to provide dorsoventral maneuverability and flexibility (Boszczyk *et al.* 2001; Jones 2015a, b) while gliding.

The cervical region was the only skeletal component measured that consistently exhibited positive allometry across all four ecotypes. Increased elongation of the cervical region with increasing size is counterintuitive in ground squirrels, since robust cervical vertebrae could provide more support for the head as skeletal size increases. In fact, across mammals, the cervical spine tends to shorten relatively with increasing body size, albeit these allometric trends differ between different clades (Arnold et al. 2017). A possible explanation for the positive cervical allometry in squirrels is that the flexibility provided by a relatively longer neck is advantageous for large squirrels regardless of ecotype, allowing more maneuverability of their necks when navigating complex terrain or burrowing. Elongation of other vertebral regions contributes to dorsal flexibility (Jones 2015a, b; Boszczyk *et al.* 2001), and elongation of the cervical region may similarly facilitate locomotion. Additionally, it is possible that an ability to see a large range of land could be advantageous for large ground squirrels. To avoid predation, large ground squirrels must seek out abundant piles of boulders and rocks for cover, while smaller ground squirrels tend to use vegetation for cover (Armstrong 2007). Larger ground squirrels’ need to scan the landscape to look for large boulders could explain selective pressures for more flexible necks. Furthermore, ground squirrels exhibit bipedal posture when threatened by potential predators (Swaisgood *et al.* 1999), and an elongate neck could better enhance the view of their environment that this posture provides, especially for large, conspicuous squirrels. Meanwhile, elongation of the cervical vertebrae could be advantageous for gliding squirrels as it could contribute to the evolution of a more elongate and aerodynamic form.

Gliding and ground squirrels exhibited a positive allometric trend for head elongation ratio. More elongate heads in larger gliders may decrease drag and make gliding more aerodynamic. The positive allometric trend in ground squirrels, however, is more surprising. Some large ground squirrel species use their heads to push stones aside when burrowing (Kwiecinski 1998), which a more robust head would better facilitate. The positive allometric trends of head elongation ratio in all ecotypes follow similar trends of craniofacial evolutionary allometry (CREA), the tendency for larger species to evolve relatively longer rostrums. CREA has been observed in a diverse range of mammalian clades, including antelopes, kangaroos, bats, and mongooses (Cardini and Polly 2013; Cardini *et al* 2015, Arbour *et al.* 2021). CREA was also observed in tree squirrels of the subfamily Sciurinae (Cardini and Polly 2013), suggesting that the elongation of the rostrum rather than the braincase contributed to the overall elongation of the cranium in squirrels.

### Predictors of body shape variation

The relative depth of the ribcage (R^2^ = 0.64) and thoracic vertebrae (R^2^ = 0.19) were the best predictors of body shape variation in squirrels, together contributing to 83% of total body shape variation. These structures are vital to supporting the body against gravity and supporting the limbs during propulsive forces (Kardong 2014). Our findings slightly differ from trends described in carnivorans, where elongation of the lumbar region (R^2^ = 0.41) followed by relative length of the ribs (R^2^ = 0.21) and elongation of the thoracic vertebrae (R^2^ = 0.14) were the best predictors of body shape variation (Law 2021b). The dorsoventral flexibility that an elongate lumbar region provides (Jones 2015a, b; Boszczyk *et al.* 2001) could allow for increased maneuverability when hunting, especially during pouncing and chasing behaviors. This could explain why the lumbar region contributes far more to body shape variation in carnivorans compared to squirrels.

We also found that, consistent with findings in other taxonomic groups, the best predictor of body shape variation differs between ecotype or clade. In squirrels, relative rib lengths explained the most variation out of any morphological component in most ecotypes (ground squirrels R^2^ = 0.34; chipmunks R^2^ = 0.48; tree squirrels R^2^ = 0.54) except for gliding squirrels, where head elongation (R^2^ = 0.55) explained the majority of body shape variation. Across carnivoran families, the best predictors of body shape variation included elongation or shortening of the head, cervical, and/or lumbar regions, as well as the relative length of the ribs (Law et al. 2019; Law 2021a). Even carnivorans that exhibit incomplete convergence towards body elongation (i.e., weasels, civets, and mongooses) display multiple pathways towards their converging elongate body plans (Law 2022). Altogether, these results demonstrate how the evolution of different body shapes can arise through multiple diverging evolutionary pathways (Ward and Mehta 2010; Wake et al. 2011; Ward and Mehta 2014; Morinaga and Bergmann 2017; Bergmann and Morinaga 2019; Law 2021a; Law 2022).

## Conclusion

Habitat use influenced allometric patterns in body shape and its underlying morphological components in Sciuridae. First, ground squirrels exhibit negative body shape allometry, while gliding squirrels and chipmunks evolved more elongate bodies with increasing size. Second, only ground squirrels exhibit a relationship between the forelimb and body shape, where more elongate species exhibit relatively shorter forelimbs. Finally, the relative length of the ribs and elongation or shortening of the thoracic region contributes the most to body shape variation across squirrels. Altogether, because body shape variation has far-reaching effects on the physiology, biomechanics, and ecology of species, including heat conservation (Brown and Lasiewski 1972), locomotion (Sharpe *et al.* 2015, Ward *et al.* 2015), and ability to exploit niches (Law 2019), these results provide a strong morphological foundation for future research investigating the evo-devo and evolutionary ecology of squirrel and other mammalian body plans.

**Fig. 6.**
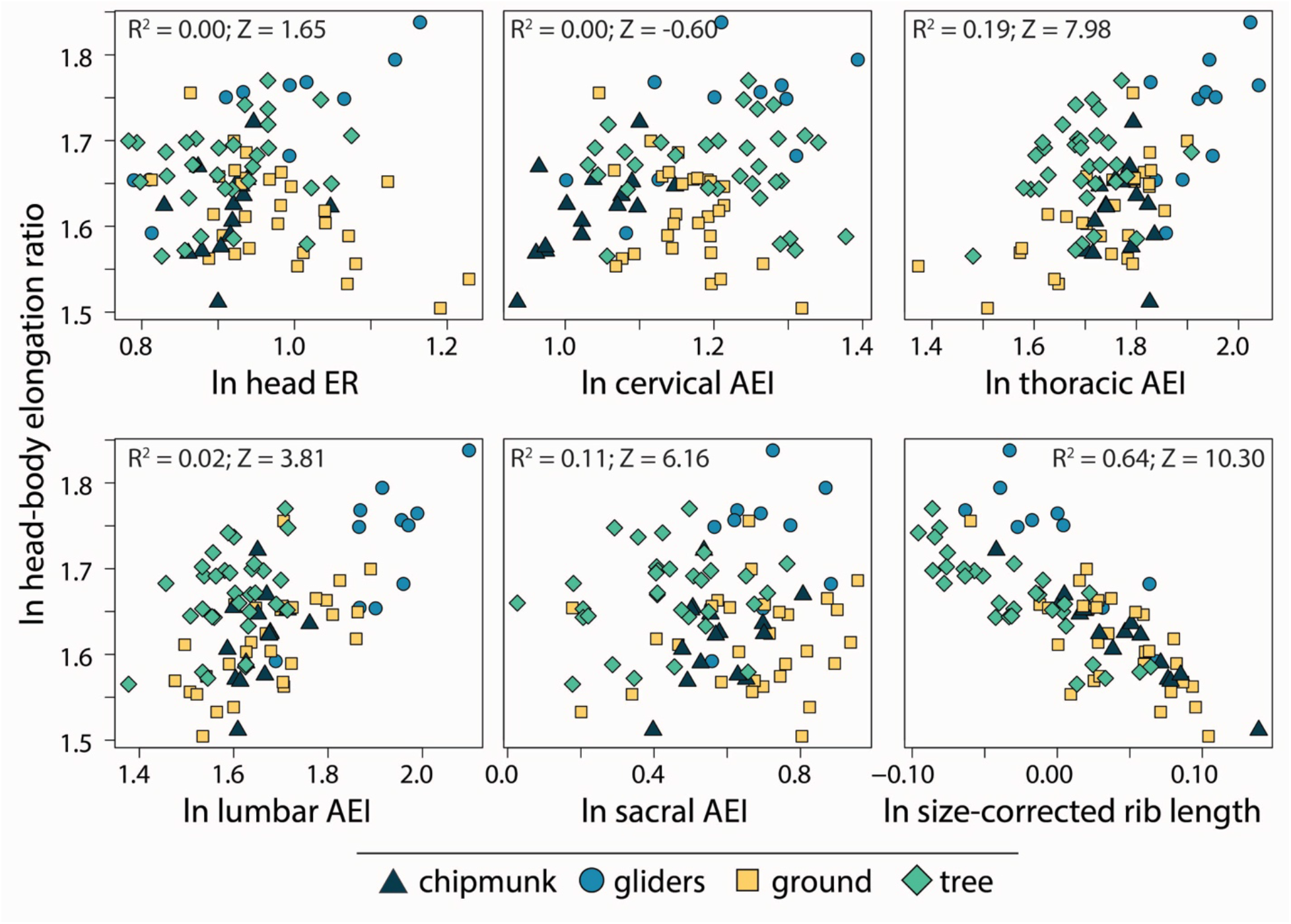
Scatterplots of head-body elongation ratio (hbER) versus skeletal components underlying body shape diversity. R^2^ and Z scores were obtained from phylogenetic multiple regression with the random residual permutation procedure (RRPP). See Table 1 for full results.

## Acknowledgements

The authors are grateful to the collections and its staff of the Beaty Biodiversity Museum at the University of British Columbia, Biodiversity Institute and Natural History Museum at Kansas University, Burke Museum of Natural History and Culture at the University of Washington, Florida Museum of Natural History, Museum of Comparative Zoology at Harvard University, Museum of Vertebrate Zoology at UC Berkeley, Natural History Museum of Los Angeles County, University of Puget Sound Museum, National Museum of Natural History. T.J.L. was supported by the American Museum of Natural History REU program; A.E.B. was supported by the American Museum of Natural History REU program, the Mary Gates Endowment, and the American Society of Mammalogists Grant-in-Aid of Research Award; J.R. and A.M. were supported by a National Science Foundation Postdoctoral Research Fellowship REU Program; S.E.S. was supported by National Science Foundation IOS-2017738; and C.J.L. was supported by National Science Foundation DBI-1906248 and DBI–2128146, the Gerstner Family Foundation and the Richard Gilder Graduate School at the American Museum of Natural History, an Iuvo Postdoctoral Award at the University of Washington, and a University of Texas Early Career Provost Fellowship.

