## Supplementary Materials for "Scaling patterns of body plans differ among squirrel ecotypes"

Table S1. Catalog numbers of specimens used

Table S2. Slope and intercept coefficients of PGLS models between body shape components, size, and ecotype

**Table S1**. Catalog numbers of specimens used. KU = Biodiversity Institute and Natural History Museum; LACM = Natural History Museum of Los Angeles County; MCZ = Museum of Comparative Zoology; MVZ = Museum of Vertebrate Zoology; PSMP = University of Puget Sound Museum; UBC = Beaty Museum; UF = Florida Museum of Natural History; USNM = National Museum of Natural History; UWBM = Burke Museum of Natural History and Culture

| species | sex | ecotype | catalog |
|---|---|---|---|
| Tamias_alpinus | F | chipmunk | MVZ207209 |
| Tamias_alpinus | F | chipmunk | MVZ207210 |
| Tamias_amoenus | F | chipmunk | UWBM31698 |
| Tamias_amoenus | M | chipmunk | UWBM35963 |
| Tamias_amoenus | F | chipmunk | UWBM39189 |
| Tamias_amoenus | F | chipmunk | UWBM75475 |
| Tamias_amoenus | F | chipmunk | UWBM78184 |
| Tamias_cinereicollis | F | chipmunk | UWBM38257 |
| Tamias_dorsalis | M | chipmunk | UWBM78903 |
| Tamias_dorsalis | F | chipmunk | UWBM78908 |
| Tamias_dorsalis | M | chipmunk | UWBM79657 |
| Tamias_merriami | M | chipmunk | UWBM60172 |
| Tamias_merriami | F | chipmunk | UWBM60174 |
| Tamias_minimus | F | chipmunk | MVZ219925 |
| Tamias_minimus | M | chipmunk | MVZ219926 |
| Tamias_minimus | M | chipmunk | UWBM30813 |
| Tamias_minimus | M | chipmunk | UWBM77677 |
| Tamias_minimus | F | chipmunk | UWBM77683 |
| Tamias_panamintinus | F | chipmunk | MVZ224274 |
| Tamias_panamintinus | M | chipmunk | MVZ224275 |
| Tamias_quadrivittatus | F | chipmunk | UWBM35298 |
| Tamias_quadrivittatus | F | chipmunk | UWBM35298 |
| Tamias_quadrivittatus | F | chipmunk | UWBM35299 |
| Tamias_quadrivittatus | F | chipmunk | UWBM35299 |
| Tamias_quadrivittatus | M | chipmunk | UWBM35301 |

| | | | |
|---|---|---|---|
| Tamias_quadrivittatus | F | chipmunk | UWBM36479 |
| Tamias_quadrivittatus | F | chipmunk | UWBM36480 |
| Tamias_ruficaudus | F | chipmunk | UWBM34354 |
| Tamias_ruficaudus | M | chipmunk | UWBM34355 |
| Tamias_ruficaudus | M | chipmunk | UWBM34357 |
| Tamias_ruficaudus | M | chipmunk | UWBM34357 |
| Tamias_ruficaudus | M | chipmunk | UWBM34360 |
| Tamias_ruficaudus | M | chipmunk | UWBM76595 |
| Tamias_senex | F | chipmunk | UWBM78783 |
| Tamias_sibiricus | F | chipmunk | UWBM39255 |
| Tamias_sibiricus | F | chipmunk | UWBM39255 |
| Tamias_sibiricus | F | chipmunk | UWBM39257 |
| Tamias_sibiricus | M | chipmunk | UWBM39258 |
| Tamias_sibiricus | F | chipmunk | UWBM39259 |
| Tamias_sibiricus | M | chipmunk | UWBM77286 |
| Tamias_siskiyou | U | chipmunk | UWBM80827 |
| Tamias_siskiyou | F | chipmunk | UWBM80828 |
| Tamias_siskiyou | F | chipmunk | UWBM80829 |
| Tamias_speciosus | F | chipmunk | UWBM43059 |
| Tamias_speciosus | F | chipmunk | UWBM60295 |
| Tamias_speciosus | F | chipmunk | UWBM60297 |
| Tamias_speciosus | M | chipmunk | UWBM60299 |
| Tamias_striatus | M | chipmunk | UWBM35246 |
| Tamias_townsendii | F | chipmunk | UWBM35167 |
| Tamias_townsendii | F | chipmunk | UWBM35180 |
| Tamias_townsendii | F | chipmunk | UWBM35182 |
| Tamias_townsendii | F | chipmunk | UWBM35208 |
| Tamias_townsendii | F | chipmunk | UWBM35213 |
| Aeretes_melanopterus | U | gliding | MCZ19994 |
| Eoglaucomys_fimbriatus | F | gliding | USNM173363 |
| Eoglaucomys_fimbriatus | F | gliding | USNM173365 |
| Eupetaurus_cinereus | M | gliding | UF28583 |
| Glaucomys_sabrinus | M | gliding | UWBM35051 |
| Glaucomys_sabrinus | F | gliding | UWBM35053 |
| Glaucomys_sabrinus | F | gliding | UWBM35056 |
| Glaucomys_sabrinus | F | gliding | UWBM35058 |
| Glaucomys_sabrinus | M | gliding | UWBM35077 |
| Glaucomys_volans | M | gliding | MVZ97094 |
| Glaucomys_volans | F | gliding | UWBM35225 |
| Glaucomys_volans | F | gliding | UWBM43897 |

| | | | |
|---|---|---|---|
| Iomys_horsfieldii | F | gliding | PSM10370 |
| Petaurista_alborufus | F | gliding | MVZ174855 |
| Petaurista_alborufus | U | gliding | MVZ183717 |
| Petaurista_elegans | U | gliding | MCZ36579 |
| Petaurista_elegans | U | gliding | MCZ36580 |
| Petaurista_petaurista | F | gliding | USNM197320 |
| Petaurista_petaurista | M | gliding | USNM588884 |
| Petaurista_philippensis | M | gliding | USNM314973 |
| Pteromys_volans | M | gliding | UWBM39689 |
| Ammospermophilus_leucurus | F | ground | MVZ216222 |
| Ammospermophilus_leucurus | F | ground | UWBM74639 |
| Ammospermophilus_leucurus | F | ground | UWBM74641 |
| Ammospermophilus_leucurus | F | ground | UWBM74643 |
| Ammospermophilus_leucurus | M | ground | UWBM74646 |
| Ammospermophilus_leucurus | M | ground | UWBM74647 |
| Ammospermophilus_nelsoni | F | ground | MVZ234368 |
| Callospermophilus_lateralis | M | ground | UWBM38959 |
| Callospermophilus_lateralis | F | ground | UWBM38960 |
| Callospermophilus_lateralis | F | ground | UWBM38961 |
| Callospermophilus_lateralis | F | ground | UWBM38964 |
| Callospermophilus_lateralis | F | ground | UWBM38965 |
| Callospermophilus_madrensis | M | ground | MVZ99799 |
| Callospermophilus_saturatus | M | ground | UWBM31137 |
| Callospermophilus_saturatus | M | ground | UWBM38984 |
| Callospermophilus_saturatus | F | ground | UWBM44578 |
| Callospermophilus_saturatus | F | ground | UWBM44586 |
| Cynomys_gunnisoni | M | ground | MVZ99763 |
| Cynomys_ludovicianus | M | ground | UWBM75774 |
| Cynomys_mexicanus | M | ground | MVZ91189 |
| Ictidomys_mexicanus | M | ground | MVZ93788 |
| Ictidomys_mexicanus | F | ground | MVZ93789 |
| Marmota_broweri | M | ground | UWBM32251 |
| Marmota_broweri | M | ground | UWBM82831 |
| Marmota_caligata | F | ground | UWBM13561 |
| Marmota_caligata | F | ground | UWBM31095 |
| Marmota_caligata | M | ground | UWBM35529 |
| Marmota_caligata | M | ground | UWBM35997 |
| Marmota_caligata | U | ground | UWBM43431 |
| Marmota_flaviventris | F | ground | UWBM32500 |
| Marmota_flaviventris | M | ground | UWBM77395 |

| | | | |
|---|---|---|---|
| Marmota_flaviventris | M | ground | UWBM82825 |
| Marmota_monax | F | ground | UWBM38327 |
| Marmota_monax | M | ground | UWBM39792 |
| Marmota_monax | F | ground | UWBM82380 |
| Marmota_olympus | F | ground | PSMP2520 |
| Marmota_vancouverensis | M | ground | UBC019561 |
| Marmota_vancouverensis | M | ground | UBC019586 |
| Otospermophilus_beecheyi | U | ground | UWBM15430 |
| Otospermophilus_beecheyi | M | ground | UWBM37043 |
| Otospermophilus_beecheyi | M | ground | UWBM39380 |
| Otospermophilus_beecheyi | F | ground | UWBM80593 |
| Otospermophilus_variegatus | M | ground | UWBM79876 |
| Poliocitellus_franklinii | F | ground | UWBM33260 |
| Poliocitellus_franklinii | F | ground | UWBM33261 |
| Urocitellus_armatus | F | ground | MVZ64641 |
| Urocitellus_armatus | F | ground | MVZ72124 |
| Urocitellus_beldingi | F | ground | UWBM42455 |
| Urocitellus_beldingi | F | ground | UWBM42456 |
| Urocitellus_beldingi | U | ground | UWBM42457 |
| Urocitellus_beldingi | F | ground | UWBM42458 |
| Urocitellus_beldingi | U | ground | UWBM42459 |
| Urocitellus_beldingi | F | ground | UWBM80581 |
| Urocitellus_columbianus | F | ground | UWBM12600 |
| Urocitellus_columbianus | M | ground | UWBM14267 |
| Urocitellus_columbianus | F | ground | UWBM37533 |
| Urocitellus_columbianus | F | ground | UWBM74770 |
| Urocitellus_columbianus | M | ground | UWBM80405 |
| Urocitellus_elegans | M | ground | UWBM33286 |
| Urocitellus_parryii | F | ground | UWBM39263 |
| Urocitellus_parryii | F | ground | UWBM39266 |
| Urocitellus_parryii | F | ground | UWBM39268 |
| Urocitellus_parryii | F | ground | UWBM39270 |
| Urocitellus_parryii | M | ground | UWBM39271 |
| Urocitellus_parryii | F | ground | UWBM39623 |
| Urocitellus_richardsonii | M | ground | UWBM32960 |
| Urocitellus_richardsonii | M | ground | UWBM32961 |
| Urocitellus_richardsonii | M | ground | UWBM32962 |
| Urocitellus_townsendii | U | ground | UWBM28071 |
| Urocitellus_townsendii | F | ground | UWBM78314 |
| Urocitellus_townsendii | F | ground | UWBM78333 |

| | | | |
|---|---|---|---|
| Xerospermophilus_spilosoma | F | ground | UWBM35279 |
| Xerospermophilus_spilosoma | F | ground | UWBM35282 |
| Xerospermophilus_spilosoma | M | ground | UWBM35284 |
| Xerospermophilus_spilosoma | M | ground | UWBM35285 |
| Xerospermophilus_spilosoma | M | ground | UWBM35286 |
| Xerospermophilus_tereticaudus | F | ground | UWBM35502 |
| Xerus_erythropus | M | ground | MVZ115437 |
| Xerus_inauris | F | ground | MVZ117287 |
| Callosciurus_erythraeus | F | tree | CAS6370 |
| Callosciurus_erythraeus | F | tree | CAS6371 |
| Callosciurus_prevostii | M | tree | LACM097638 |
| Callosciurus_prevostii | F | tree | LACM97346 |
| Callosciurus_pygerythrus | F | tree | PSM16608 |
| Callosciurus_pygerythrus | M | tree | PSM27419 |
| Funambulus_palmarum | F | tree | MVZ183704 |
| Funambulus_palmarum | M | tree | MVZ183706 |
| Funambulus_pennantii | F | tree | UF28826 |
| Funambulus_pennantii | M | tree | UF30290 |
| Funisciurus_pyrropus | F | tree | MVZ196233 |
| Heliosciurus_mutabilis | M | tree | MVZ220919 |
| Heliosciurus_rufobrachium | U | tree | MCZ35324 |
| Heliosciurus_rufobrachium | U | tree | MCZ35327 |
| Heliosciurus_rufobrachium | M | tree | MVZ196227 |
| Microsciurus_flaviventer | M | tree | MVZ124046 |
| Microsciurus_flaviventer | M | tree | MVZ154935 |
| Microsciurus_flaviventer | F | tree | MVZ154936 |
| Paraxerus_boehmi | F | tree | LSUMZ37843 |
| Prosciurillus_murinus | M | tree | LSUMZ38363 |
| Protoxerus_stangeri | F | tree | MVZ196228 |
| Protoxerus_stangeri | M | tree | MVZ196230 |
| Ratufa_bicolor | M | tree | MVZ123699 |
| Ratufa_macroura | U | tree | UWBM21183 |
| Sciurus_aberti | M | tree | UWBM35277 |
| Sciurus_aberti | F | tree | UWBM35278 |
| Sciurus_aberti | F | tree | UWBM74145 |
| Sciurus_aestuans | U | tree | MVZ182069 |
| Sciurus_aestuans | F | tree | MVZ182070 |
| Sciurus_aureogaster | F | tree | LACM053653 |
| Sciurus_aureogaster | F | tree | LACM053654 |
| Sciurus_carolinensis | F | tree | UWBM35488 |

| | | | |
|---|---|---|---|
| Sciurus_carolinensis | M | tree | UWBM39373 |
| Sciurus_carolinensis | M | tree | UWBM39787 |
| Sciurus_carolinensis | F | tree | UWBM43904 |
| Sciurus_carolinensis | F | tree | UWBM75810 |
| Sciurus_colliaei | F | tree | LACM058790 |
| Sciurus_deppei | M | tree | MVZ98319 |
| Sciurus_deppei | F | tree | MVZ98326 |
| Sciurus_griseus | F | tree | UWBM35487 |
| Sciurus_griseus | M | tree | UWBM42330 |
| Sciurus_griseus | F | tree | UWBM74097 |
| Sciurus_griseus | F | tree | UWBM76257 |
| Sciurus_griseus | F | tree | UWBM76264 |
| Sciurus_nayaritensis | M | tree | MVZ109632 |
| Sciurus_niger | M | tree | UWBM35226 |
| Sciurus_niger | F | tree | UWBM35229 |
| Sciurus_niger | F | tree | UWBM35230 |
| Sciurus_niger | M | tree | UWBM35231 |
| Sciurus_niger | M | tree | UWBM81529 |
| Sciurus_spadiceus | U | tree | MVZ166023 |
| Sciurus_stramineus | M | tree | MVZ135638 |
| Sciurus_variegatoides | M | tree | MVZ131091 |
| Sciurus_variegatoides | F | tree | MVZ131095 |
| Sciurus_vulgaris | M | tree | UWBM39069 |
| Sciurus_vulgaris | F | tree | UWBM39260 |
| Sciurus_vulgaris | F | tree | UWBM77295 |
| Sundasciurus_juvencus | U | tree | KU165382 |
| Tamiasciurus_douglasii | F | tree | UWBM39137 |
| Tamiasciurus_douglasii | F | tree | UWBM44366 |
| Tamiasciurus_douglasii | F | tree | UWBM81947 |
| Tamiasciurus_douglasii | F | tree | UWBM82060 |
| Tamiasciurus_douglasii | F | tree | UWBM82088 |
| Tamiasciurus_hudsonicus | M | tree | UWBM35237 |
| Tamiasciurus_hudsonicus | F | tree | UWBM81502 |
| Tamiasciurus_hudsonicus | M | tree | UWBM81510 |
| Tamiasciurus_hudsonicus | M | tree | UWBM81909 |
| Tamiasciurus_hudsonicus | F | tree | UWBM82174 |
| Tamiops_maritimus | M | tree | MVZ186481 |
| Tamiops_swinhoei | F | tree | UWBM75283 |

**Table S2**. Slope and intercept coefficients of PGLS models between body shape components, size, and ecotype. 95% bootstrap confidence intervals were used to determine if body size-shape relationships were allometric. Bolded values indicate slopes deviated from isometry.

| ecotype | intercept | intercept L95% | intercept U95% | Slop e | Slope L95% | Slope U95% |
|---|---|---|---|---|---|---|
| ln head-body elongation ratio | | | | | | |
| all squirrels | 1.73 | 1.56 | 1.9 | -0.02 | -0.07 | 0.02 |
| chipmunk | 0.9 | 0.27 | 1.58 | **0.24** | **0.01** | **0.45** |
| gliding | 1.27 | 1.03 | 1.5 | **0.12** | **0.06** | **0.19** |
| ground | 2.01 | 1.83 | 2.17 | **-0.11** | **-0.15** | **-0.06** |
| tree | 1.74 | 1.55 | 1.94 | -0.02 | -0.08 | 0.03 |
| ln head elongation ratio | | | | | | |
| all squirrels | 0.31 | 0.11 | 0.49 | **0.18** | **0.13** | **0.23** |
| chipmunk | 0.27 | -0.59 | 1.11 | 0.21 | -0.06 | 0.5 |
| gliding | 0.19 | -0.14 | 0.5 | **0.21** | **0.13** | **0.3** |
| ground | 0.17 | -0.07 | 0.38 | **0.22** | **0.16** | **0.29** |
| tree | 0.7 | 0.43 | 0.95 | 0.06 | -0.01 | 0.14 |
| ln cervical AEI | | | | | | |
| all squirrels | 0.4 | 0.22 | 0.56 | **0.22** | **0.18** | **0.26** |
| chipmunk | -0.45 | -1.06 | 0.1 | **0.5** | **0.32** | **0.69** |
| gliding | 0.14 | -0.16 | 0.46 | **0.29** | **0.2** | **0.37** |
| ground | 0.69 | 0.49 | 0.91 | **0.13** | **0.07** | **0.18** |
| tree | 0.23 | -0.03 | 0.49 | **0.27** | **0.2** | **0.34** |
| ln thoracic AEI | | | | | | |
| all squirrels | 1.94 | 1.65 | 2.21 | -0.06 | -0.13 | 0.02 |
| chipmunk | 1.72 | 0.85 | 2.68 | 0.01 | -0.3 | 0.29 |
| gliding | 1.46 | 0.97 | 1.97 | 0.12 | -0.01 | 0.26 |
| ground | 2.26 | 1.94 | 2.61 | **-0.17** | **-0.26** | **-0.09** |
| tree | 1.66 | 1.21 | 2.1 | 0.01 | -0.11 | 0.13 |
| ln lumbar AEI | | | | | | |
| all squirrels | 1.97 | 1.68 | 2.27 | **-0.09** | **-0.16** | **-0.02** |
| chipmunk | 0.98 | 0.19 | 1.88 | 0.21 | -0.08 | 0.48 |
| gliding | 1.36 | 0.87 | 1.91 | **0.15** | **0** | **0.28** |
| ground | 2.44 | 2.11 | 2.78 | **-0.23** | **-0.31** | **-0.15** |
| tree | 1.73 | 1.3 | 2.12 | -0.04 | -0.15 | 0.08 |

| | | | | | | |
|---|---|---|---|---|---|---|
| ln sacral AEI | | | | | | |
| all squirrels | 0.7 | 0.38 | 1.01 | -0.01 | -0.09 | 0.08 |
| chipmunk | 0.45 | -0.98 | 1.88 | 0.09 | -0.39 | 0.56 |
| gliding | 0.35 | -0.24 | 0.95 | 0.12 | -0.04 | 0.28 |
| ground | 0.82 | 0.41 | 1.17 | -0.01 | -0.11 | 0.1 |
| tree | 0.57 | 0.16 | 0.99 | 0.01 | -0.11 | 0.13 |
| ln size-corrected rib length | | | | | | |
| all squirrels | 0 | -0.12 | 0.12 | 0 | -0.03 | 0.03 |
| chipmunk | 0.57 | 0.02 | 1.09 | -0.17 | -0.35 | 0.01 |
| gliding | 0.13 | -0.07 | 0.34 | -0.03 | -0.09 | 0.02 |
| ground | -0.12 | -0.27 | 0.02 | **0.04** | **0.01** | **0.09** |
| tree | -0.02 | -0.18 | 0.13 | 0 | -0.04 | 0.04 |

**Table S3**. Slope and intercept coefficients of PGLS models between limb length and body shape. 95% bootstrap confidence intervals were used to determine if body size-shape relationships were allometric. Bolded values indicate slopes deviated from isometry.

| ecotype | intercept | intercept L95% | intercept U95% | Slope | Slope L95% | Slope U95% |
|---|---|---|---|---|---|---|
| forelimb_long vs body shape | | | | | | |
| all squirrels | -0.13 | -0.56 | 0.28 | 0.08 | -0.17 | 0.34 |
| chipmunk | -0.08 | -0.99 | 0.82 | -0.01 | -0.57 | 0.56 |
| gliding | -0.33 | -1.11 | 0.42 | 0.37 | -0.07 | 0.82 |
| ground | 0.49 | 0.07 | 0.92 | **-0.39** | **-0.66** | **-0.13** |
| tree | -0.02 | -0.5 | 0.52 | 0.01 | -0.33 | 0.29 |
| hind limb_long vs body shape | | | | | | |
| all squirrels | -0.13 | -0.5 | 0.24 | 0.08 | -0.14 | 0.29 |
| chipmunk | 0.01 | -0.63 | 0.69 | -0.05 | -0.46 | 0.34 |
| gliding | -0.35 | -1.41 | 0.74 | 0.33 | -0.31 | 0.94 |
| ground | -0.14 | -0.65 | 0.39 | 0.02 | -0.3 | 0.34 |
| tree | 0.21 | -0.43 | 0.86 | -0.14 | -0.54 | 0.25 |
| forelimb_full vs body shape | | | | | | |
| all squirrels | 0.05 | -0.29 | 0.44 | -0.03 | -0.27 | 0.17 |
| chipmunk | -0.04 | -0.74 | 0.68 | -0.02 | -0.47 | 0.42 |
| gliding | -0.06 | -0.8 | 0.72 | 0.15 | -0.31 | 0.59 |
| ground | 0.5 | 0.12 | 0.91 | **-0.36** | **-0.62** | **-0.13** |

| | | | | | | |
|---|---|---|---|---|---|---|
| tree | 0.11 | -0.47 | 0.66 | -0.06 | -0.39 | 0.29 |
| hind limb_full vs body shape | | | | | | |
| all squirrels | -0.06 | -0.46 | 0.33 | 0.04 | -0.21 | 0.27 |
| chipmunk | 0.14 | -0.55 | 0.92 | -0.11 | -0.59 | 0.29 |
| gliding | -1.27 | -2.71 | 0.07 | **0.85** | **0.04** | **1.71** |
| ground | 0.08 | -0.46 | 0.65 | **-0.11** | **-0.46** | **0.22** |
| tree | -0.02 | -0.78 | 0.73 | 0.01 | -0.43 | 0.49 |
